# Library of model implementations for sharing deep-learning image segmentation and outcomes models

**DOI:** 10.1101/773929

**Authors:** Aditya P. Apte, Aditi Iyer, Maria Thor, Rutu Pandya, Rabia Haq, Amita Shukla-Dave, Hu Yu-Chi, Sharif Elguindi, Harini Veeraraghavan, Jung Hun Oh, Andy Jackson, Joseph O. Deasy

**Affiliations:** Department of Medical Physics, Memorial Sloan Kettering Cancer Center, New York, NY 10065 USA

**Keywords:** image segmentation, deep-learning, radiomics, radiotherapy outcomes, normal tissue complication, tumor control, model implementations, library

## Abstract

An open-source library of implementations for deep-learning based image segmentation and outcomes models is presented in this work. As oncology treatment planning becomes increasingly driven by automation, such a library of model implementations is crucial to (i) validate existing models on datasets collected at different institutions, (ii) automate segmentation, (iii) create ensembles for improving performance and (iv) incorporate validated models in the clinical workflow. The library was developed with Computational Environment for Radiological Research (CERR) software platform. CERR is a natural choice to centralize model implementations due to its comprehensiveness, popularity, and ease of use. CERR provides well-validated feature extraction for radiotherapy dosimetry and radiomics with fine control over the calculation settings. This allows users to select the appropriate feature calculation used in the model derivation. Models for automatic image segmentation are distributed via Singularity containers, with seamless i/o to and from CERR. Singularity containers allow for segmentation models to be deployed with a variety of scientific computing architectures. Deployment of models is driven by JSON configuration file, making it convenient to plug-in models. Models from the library can be called programmatically for batch evaluation. The library includes implementations for popular radiotherapy models outlined in the Quantitative Analysis of Normal Tissue Effects in the Clinic effort and recently published literature. Radiomics models include features from Image Biomarker Standardization features found to be important across multiple sites and image modalities. Deep learning-based image segmentation models include state of the art networks such as Deeplab and other problem-specific architectures. The library is distributed as GNU-copyrighted software at https://www.github.com/cerr/CERR.

## Introduction

Oncology is increasingly driven by automation and prognosis. In radiotherapy treatment planning automation is crucial for segmenting lesions and organs at risk. Segmentation along with images and radiotherapy plans are fed into prediction models to estimate prognosis resulting from the planned treatment. The treatment, in turn, can then be tailored based on the estimated prognosis. Software libraries for deriving radiomics and radiotherapy features as well as outcomes models have grown rapidly. Similarly, automatic image segmentation has seen a great deal of recent progress with deep learning. Segmentation and prognostic models are affected by the choice of feature calculation engines such as deep learning framework and their associated settings. This makes it cumbersome to use the published signatures on new datasets. This work addresses the issue of variability in model implementations by creating an open-source, validated and comprehensive library of image-segmentation, radiotherapy and radiomics-based outcomes model implementations. The library of model implementations developed in this work will be useful to (i) validate existing models on datasets collected at different institutions, (ii) automate segmentation making it less prone to inter-observer variability, (iii) create ensembles from various models in the library and in-turn improving model performance and (iv) simplify incorporating validated models in clinical workflow.

There have been various efforts to deploy implementations of prognostic models. The focus has mostly been on developing online tools useful to patients and physicians. These implementations are limited to interactive use and cannot be used in batch evaluations on datasets acquired at multiple institutions. For example, DNAmito https://predictcancer.org provides online tools and an app that supports 25 cancer models for different sites. These models use patient and radiotherapy treatment characteristics. National Cancer Registration and Analysis Service [1] https://breast.predict.nhs.uk/ provides an online tool to predict survival rate for early invasive breast cancer treatment. Memorial Sloan Kettering Cancer Center [2, 3] https://www.mskcc.org/nomograms provides online tools to predict cancer outcomes based on patient characteristics and their disease. There has been a lot of recent progress in the development of toolkits for storing and deploying deep learning segmentation models. These include traditional multi-atlas-based segmentation as well as state of the art deep learning-based models. DeepInfer [4] currently provides segmentation models for Prostate gland, biopsy needle trajectory and tip and brain white matter hyperintensities. DeepInfer can also be invoked from Slicer. NiftyNET [5], a consortium of research organizations, is TensorFlow-based open-source convolutional neural networks (CNN) platform. It provides the implementation of networks such as HighRes3DNet, 3D U-net, V-net and DeepMedic which can be used to train new models as well as share pre-trained models. Plastimatch [6] is an open-source software primarily used for image registration. It provides a module for multi-atlas segmentation. Computational Environment for Radiological Research [7] software platform has been widely used for outcomes modeling and prototyping algorithms useful in radiotherapy treatment planning. In this work, we develop a research friendly library of model implementation which is comprehensive in terms of a variety of models, viz. image segmentation, radiotherapy, and radiomics. This leads to a completely automated and reproducible pipeline from DICOM to estimated prognosis.

## Methods

The library of model implementations was developed as a module in CERR; allowing users to readily access and plug-in models. The library supports the following three classes of models: (1) Segmentation models based on deep learning, (2) Radiomics models based on imaging biomarkers, and (3) Radiotherapy models based on dose-volume histograms and cell survival. The library builds on CERR’s data import/export, visualization and standardization tools for outcomes modeling. The following sections provide details about the architecture for deploying and using these three classes of models.

### Image Segmentation models

Open-source frameworks such as Tensorflow and PyTorch are popularly used to develop deep-learning-based image segmentation models. These frameworks require the installation of various operating system-specific software dependencies. Singularity container technology was used to distribute deep learning segmentation models which allow users to securely bundle libraries such that they are compatible with a variety of scientific computing architectures. Singularity container technology presents various advantages including (i) checksums for making the software stacks reproducible (ii) compatibility with HPC systems and enterprise architectures (iii) software and data controls compliance for HIPAA and (iv) can be run by users without root privileges. Separate containers are distributed for CPU and GPU implementations. Figure 1 shows schematic of invoking deep learning segmentation models through Singularity containers.

The segmentation models often require preprocessing of input images as well as post-processing of output segmentation. These pre and post-processing options are defined through model-specific configuration files. An important pre-processing option is the selection of the region of interest used for segmentation. This must correspond to the region of interest used while deriving the model. The options for selection of such region of interest include (i) Cropping to structure bounds, (ii) Crop by a fixed amount from center, (iii) Neck crop. Figure 2 shows examples of selecting regions of interest as inputs to segmentation models. Hierarchical Data Format (HDF) is used for I/O between CERR and the segmentation container. Segmentation models derived on 2-D, 3-D and 2.5-D input images are supported. Various pre-processing filters are supported to populate different channels used by CNNs.

**Figure 2:**
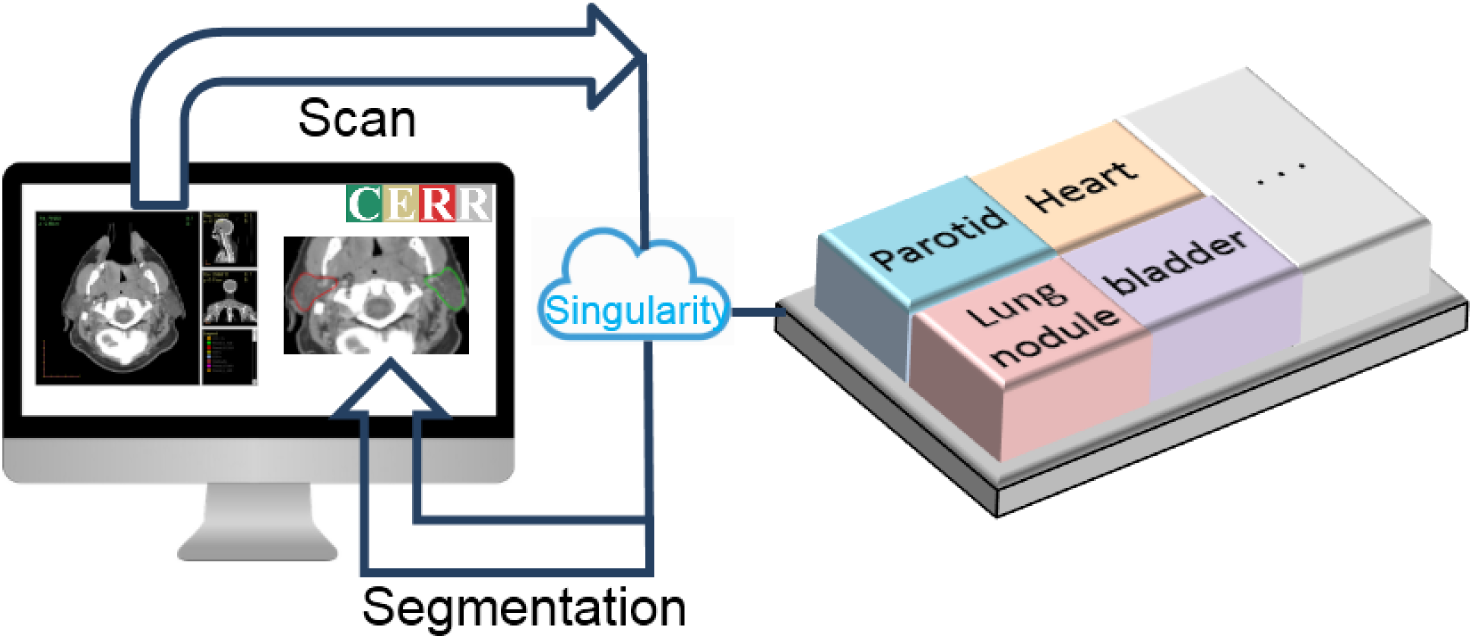
Schematic of invoking deep-learning models from CERR.

**Figure 2:**
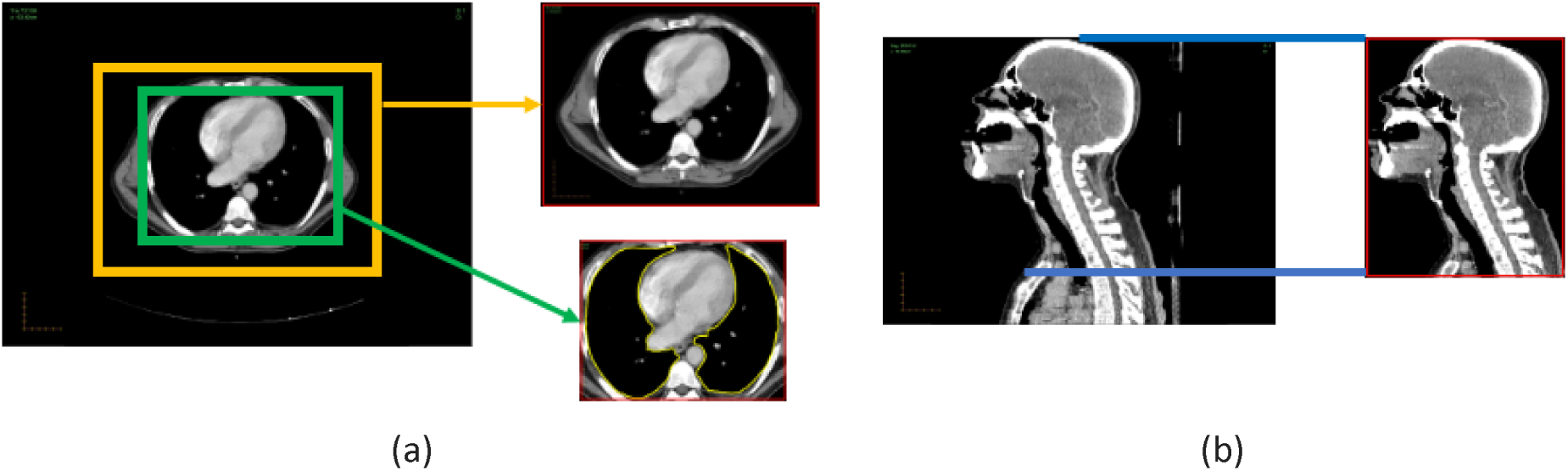
Options to select region of interest as an input to segmentation model.

The models and their parameters can be selected through the graphical user interface as shown in Figure 3. Parameters such as structure names to select region of interest can be defined using the interface.

**Figure 3:**
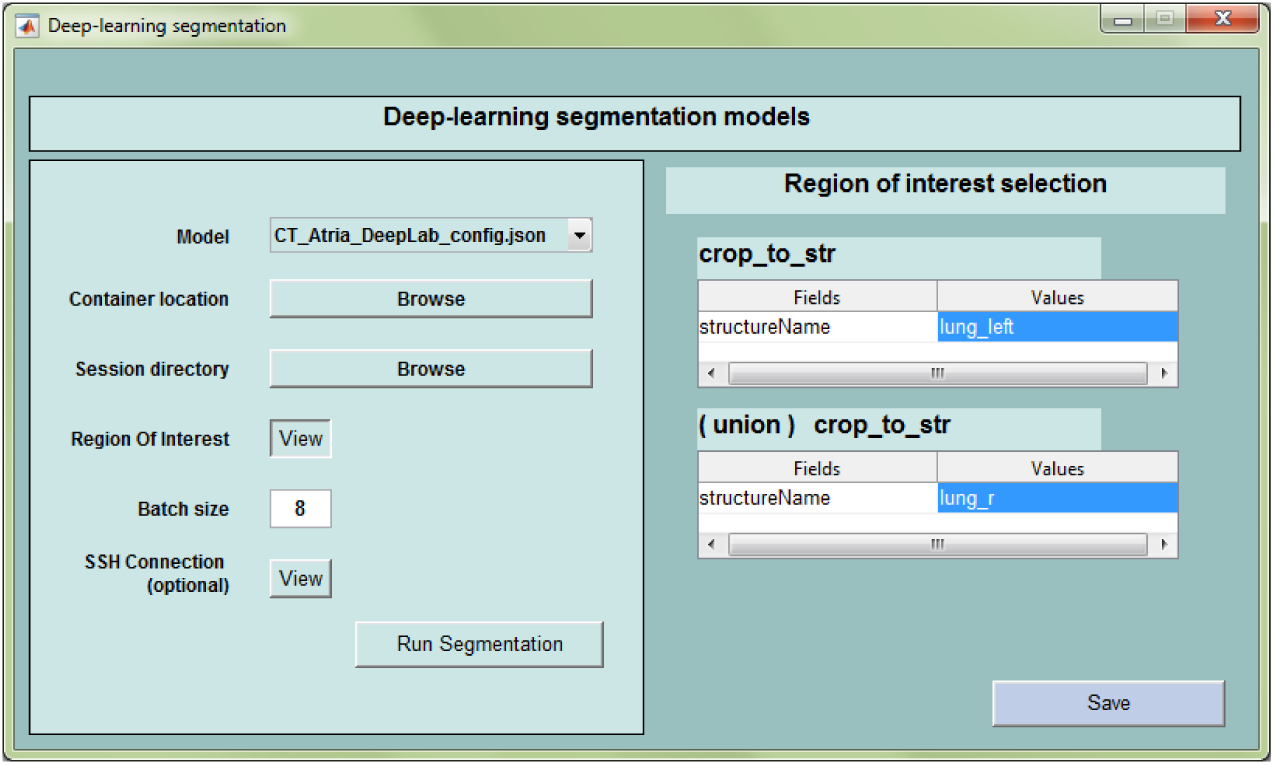
Graphical User Interface to invoke deep learning segmentation models from CERR Viewer

In addition to deep-learning-based segmentation models, multi-atlas segmentation is supported. Atlas fusion methods such as majority vote and STAPLE are supported. Plastimatch is used for deformable image registration allowing users to choose from a variety of cost functions, optimizers, and registration algorithms. This allows users to easily compare state-of-the-art with traditional segmentation models. CERR also provides metrics to evaluate the performance of segmentation such as DICE, Hausdorff distance, and deviance.

### Radiotherapy models

Radiotherapy models involve modeling Normal Tissue Complication (NTCP) and Tumor Control probabilities (TCP). NTCP models are usually based on dose-volume histogram (DVH) characteristics whereas TCP models involve estimating cell survival from radiobiology. QUANTEC series of papers provides a nice description of models for various treatment sites. CERR provides utility functions to import RT-DOSE and RT-PLAN objects as well as tools to apply linear-quadratic (LQ) corrections [8] for fraction size and to sum dose distributions from multiple courses of radiation. The library of radiotherapy models has access to the wide variety of DVH metrics in CERR such as minimum dose to hottest x percent volume (Dx), volume receiving at least x Gy of dose (Vx), mean of hottest x% volume (MOHx), mean of coldest x% volume (MOCx) and generalized equivalent dose (gEUD). Various types of NTCP models include Lyman-Kutcher-Burman (LKB) [9], linear, logistic, bi-exponential and the ability to correct for risk factors based on Appelt et al. The TCP models include cell survival model based on the dominant lesion for Prostate [10] and Jeong et al’s DVH-based model for Lung [11]. The models are defined via a JSON file as shown in figure 4.

**Figure 4:**
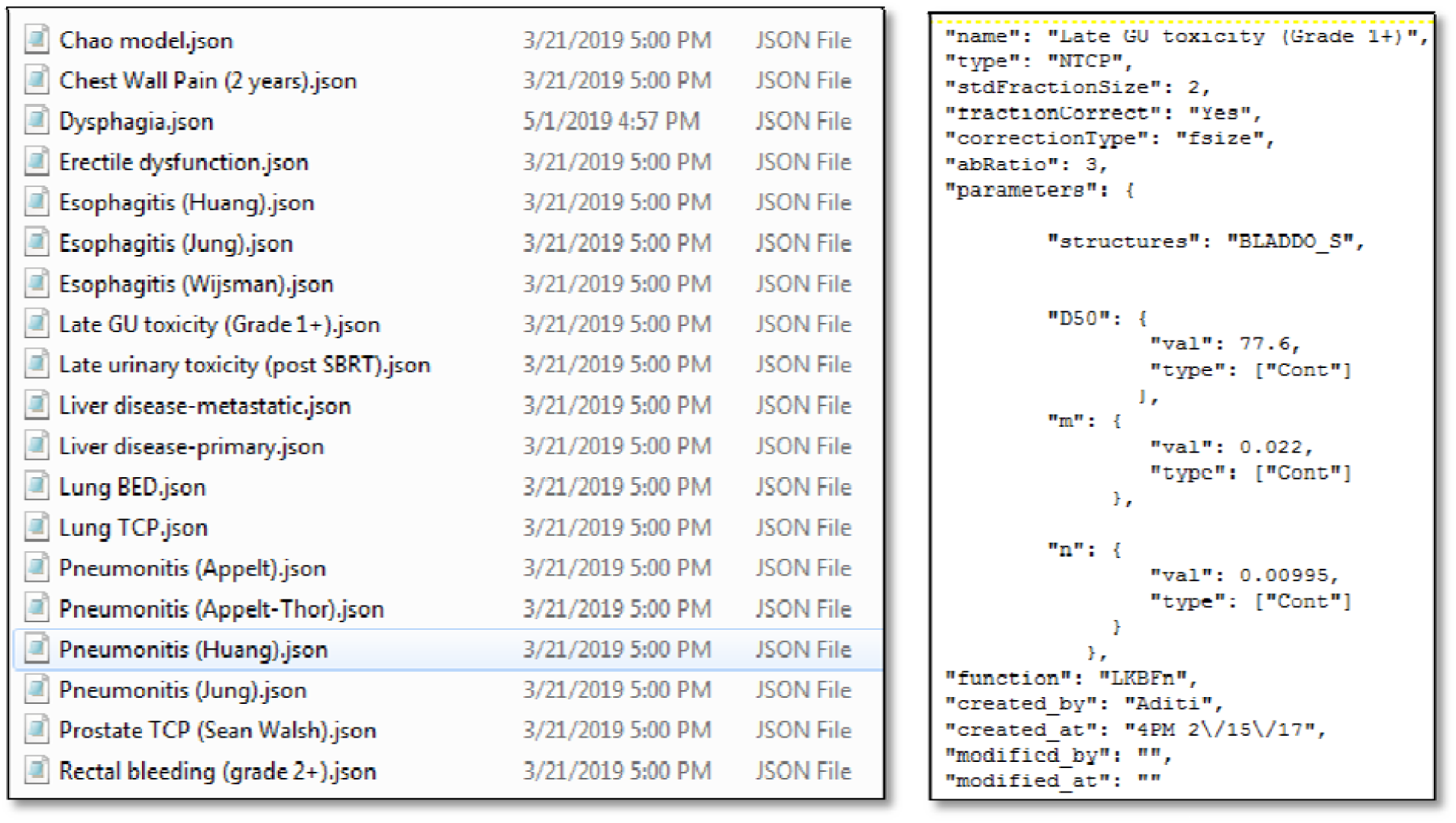
JSON files to define models.

### Radiomics models

Radiomics models require derivation of image and morphological features. These include image histogram-based features as well as texture derived from gray-level co-occurrences, run-lengths, size-zones, gray level dependence, and gray tone difference matrices. CERR supports the calculation of a variety of flavors of these features using the radiomics extensions [12]. CERR data structure supports longitudinal images as well as multiple modalities such as CT, MR, PET, SPECT, and US. This allows for the deployment of models derived with multiple modalities and various time points. For example, Hypoxia prediction model from Crispen-Ortuzar et al which used features from PET and CT scans. In addition to general radiomics features, CERR supports computation of modality-specific features such as Tofts model [13] parameters for DynamiContrast-Enhanceded (DCE) MRI and Non-model-based parameters such as time to half peak (TTHP) [14][Lee at al]. Diffusion Weighted Imaging (DWI) model parameters are supported as well. Parameter extraction is driven via configuration files in json format.

### Quality assurance

Implementations of models were validated by comparing results with those provided by the developers of these models. The results from the developers are available in the form publications, test datasets or open source code. The DVH-based features used in radiotherapy models have been independently validated by comparing with commercial treatment planning software. Radiotherapy models have been thoroughly validated by comparing with hand calculations as well as results reported in their respective publications. Radiomics features in CERR are IBSI compliant and validated with other popular libraries such as ITK and PyRadiomics [15]. Test suite for model implementation was developed using test datasets. This ensures accurate implementation and reproducibility with respect to software updates. The Test suite along with test datasets are distributed along with CERR.

## Results

CERR’s Model Implementation Library is distributed as an open-source, GNU copyrighted software with the CERR platform. Documentation for the available models and their usage is available athttps://github.com/cerr/CERR/wiki. Tables 1-3 list various models implemented for image segmentation, radiotherapy, and radiomics based outcomes prediction. Image segmentation models include the popular Deeplab [16] network for segmentation of prostate, heart sub structures, chewing and swallowing structures as well as specialized architectures such as MRRN [17] for segmentation of lung nodules. Radiotherapy and radiomics models derived with a sufficient number of patients as well as satisfactory performance on validation dataset were included. Radiotherapy based models include the prediction of normal tissue complication resulting from Lung and Prostate radiotherapy. Radiomics models include features that have been shown to be predictive in a variety of studies. For example, the features used in overall survival prediction model for lung cancer patients have been shown to be important across multiple treatment sites and image types. Similarly, the ColLage features have been shown to predict brain necrosis as well as breast cancer sub-types. Segmentation model for cardiac sub-structures [Haq et al] was derived at Memorial Sloan Kettering Cancer Center and used by the University of Pennsylvania to segment over 400 scans from the RTOG 0617 dataset. Features extracted from these sub-structures will be used for outcomes modeling.

**Table 1.**
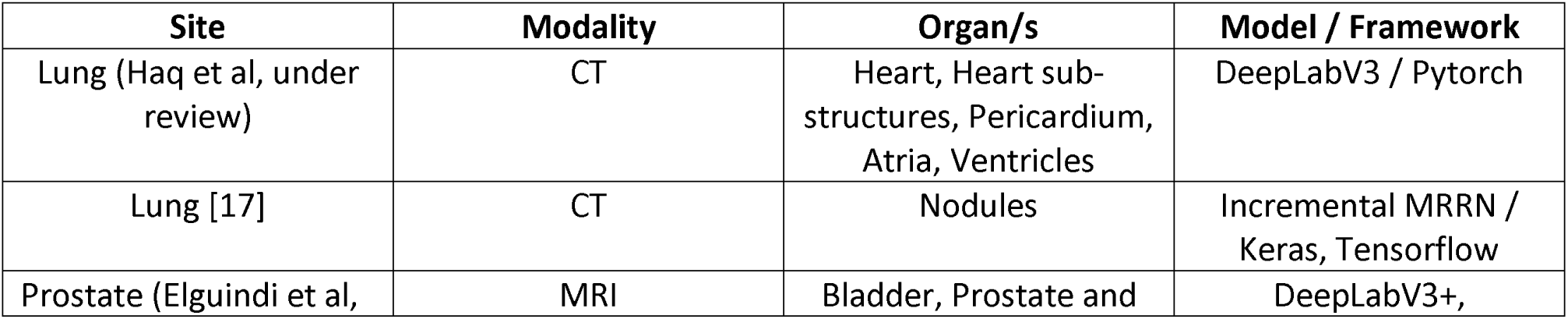
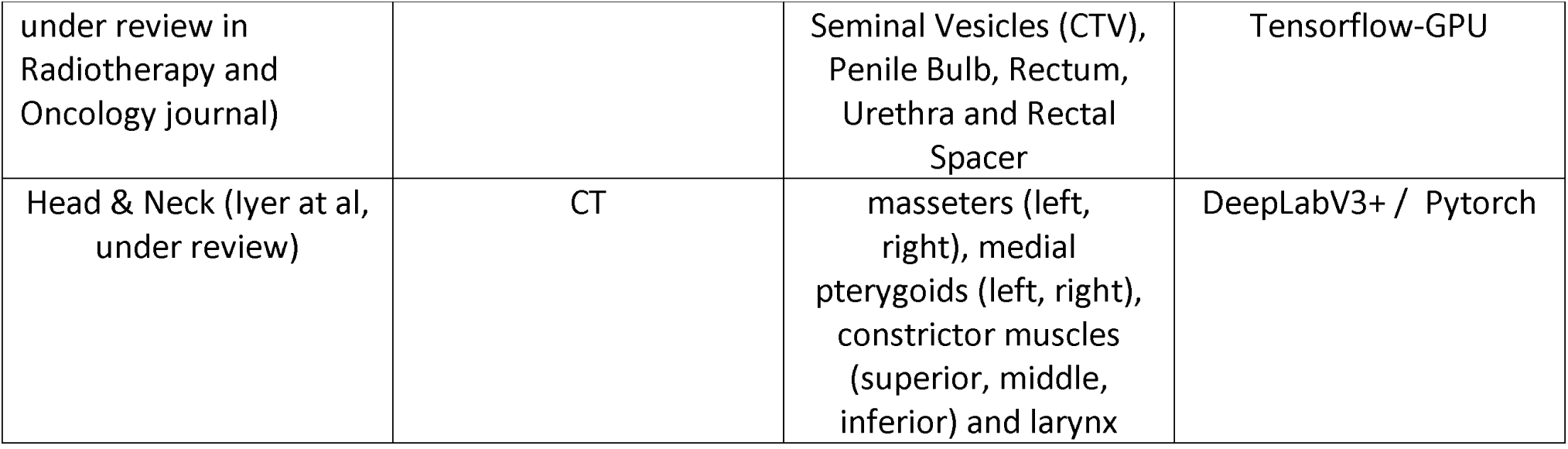
Image segmentation models.

**Table 2.** Radiotherapy models.

| Site | Endpoint | Model |
| --- | --- | --- |
| Prostate [18-21] | Rectal bleeding | LKB |
|  | Late urinary toxicity |  |
|  | Late urinary toxicity post SBRT |  |
| Lung[22] | Radiation-induced liver disease | LKB |
| Prostate | Erectile dysfunction | Linear regression |
| Lung [23] | Esophagitis | Logistic regression |
| Lung[24] | Dysphagia | Logistic regression |
| Head & Neck[25] | Xerostomia | Bi-exponential |
| Lung[26] | Chest wall pain | Non-standard |
| Lung[27] | Pneumonitis | Logistic with correction for clinical risk factors |
| Lung[10] | Tumor control probability | Radiobiological |
| Prostate[28] | Tumor control probability | Logistic |

**Table 3.** Radiotherapy models.

| Site | Endpoint | Features |
| --- | --- | --- |
| Ovarian [29] | Survival for high grade Ovarian cancer | CluDiss (inter-tumor heterogeneity) |
| Breast, Brain [30] | (i) Radiation necrosis in T1-w MRI Brain, (ii) Subtype classification in DCE-MRI Breast | CoLLAGe (gradient orientations) |
| Head & Neck [31] | Hypoxia | P90 of FDG PET SUV, LRHGLE of CT |
| Lung, H&N [32] | Survival for NSCLC and HNSCC | Statistics Energy, Shape Compactness, Grey Level Nonuniformity, wavelet Grey Level Nonuniformity HLH |

## Discussion

In addition to validating models, the library of models can be used to create ensembles for improving performance. The library is useful in inter-institutional validation as well as collaboration. The library of models has been used to build tools useful to personalize treatment planning and to drive clinical segmentation. The following describes the applications that make use of the library of model implementations.

### Radiotherapy Outcomes Estimator

Radiotherapy Outcomes Estimator (ROE) is a tool for exploring the impact of dose scaling on Tumor Control Probability (TCP) and Normal Tissue Complication Probability (NTCP) in radiotherapy. The Users define outcomes models and clinical constraints, following a simple pre-defined syntax. Inputs are specified in easily edited JavaScript Object Notation (JSON) treatment protocol files. Each file specifies tumor and critical structures; associated outcome models; model parameters including clinical risk factors (age, gender, stage etc.); and clinical constraints on outcomes. The predictive models include Lyman-Kutcher-Burman [], linear, biexponential, logistic, and the TCP model of Walsh et al. The tool simultaneously displays the TCP/NTCP dose responses as functions on the prescription dose and indicates the dose scale where the clinical limits are violated. Users can modify the clinical and model variables to visualize the simultaneous impact on TCP/NTCP. Figure 5 shows the ROE graphical user interface.

**Figure 5:**
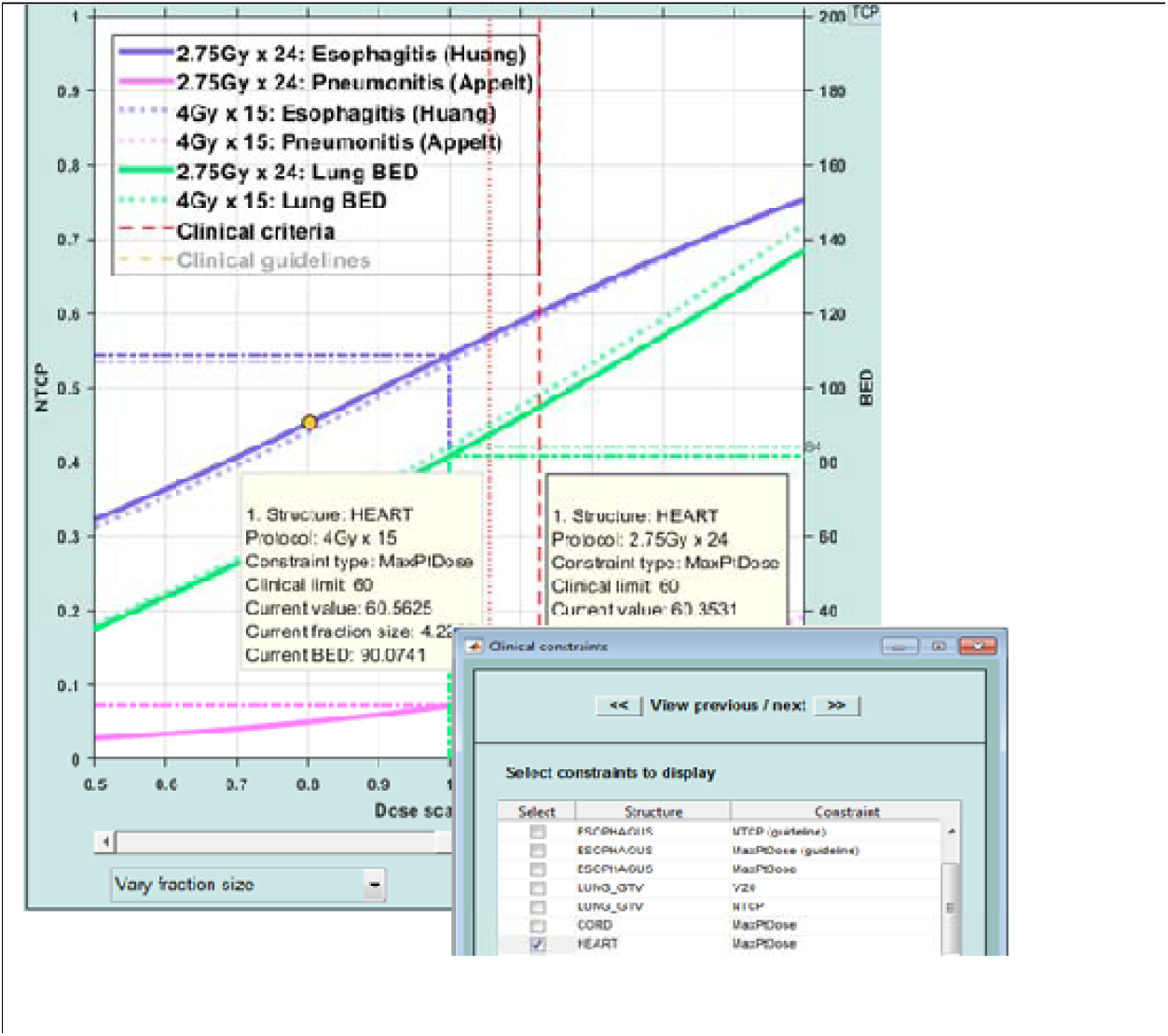
Radiotherapy Outcomes Explorer (ROE). NTCP curves for esophagitis and pneumonitis models and TCP curve for BED are shown in this example. The user interface shows points on the NTCP curves where the clinical limits are reached.

### Ensemble Voxel Attributor

Ensemble Voxel Attributor (EVA) is an extensible pipeline developed at Memorial Sloan Kettering Cancer Center to deploy deep learning segmentation algorithms into the clinic. It works in conjunction with MIM software and uses segmentation models from CERR’s library. EVA has been deployed in clinical use for segmentation of Prostate structures [Elguindi et al] and is currently undergoing testing for other sites. Figure 6 shows the workflow for EVA. The models from the library are invoked on the processing server and resulting segmentation is archived in MIM Software.

**Figure 6:**
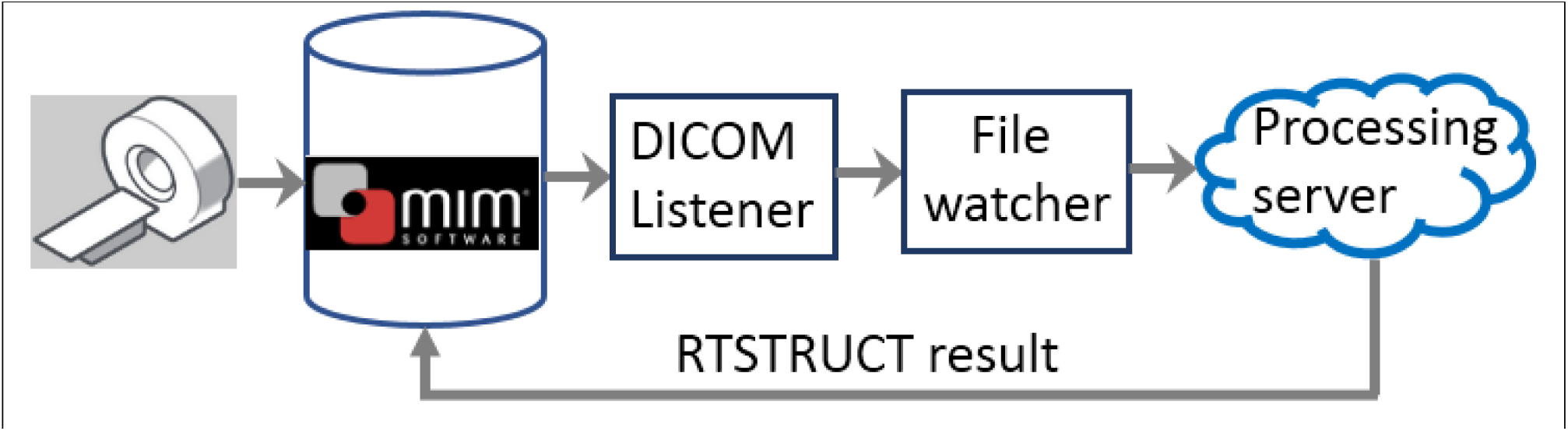
schematic of EVA pipeline.

## Conclusion

The library of model implementations for segmentation and outcomes models was developed with the CERR platform. The library makes it efficient to use and deploy deep learning segmentation, radiotherapy and radiomics based models. It leads to reproducible application of models and automated pipelines starting from imaging and radiotherapy input to the final model. This was demonstrated by sharing segmentation model for cardiac sub-structures between two institutions.

## Acknowledgements

This research was partially funded by NIH grant 1R01CA198121 and NIH/NCI Cancer Center Support grant P30 CA008748.

